# A Plant-Specific Polarity Module Establishes Cell Fate Asymmetry in the Arabidopsis Stomatal Lineage

**DOI:** 10.1101/614636

**Authors:** Matthew H. Rowe, Juan Dong, Annika K. Weimer, Dominique C. Bergmann

**Affiliations:** HHMI, Stanford University, Stanford, CA, USA 94305-5020; Department of Biology, Stanford University, Stanford, CA, USA 94305-5020; Waksman Institute of Microbiology, Rutgers, the State University of New Jersey, Piscataway, NJ 08901

## Abstract

Generating cell polarity in anticipation of asymmetric cell division is required in many developmental contexts across a diverse range of species. Physical and genetic diversity among major multicellular taxa, however, demand different molecular solutions to this problem. The Arabidopsis stomatal lineage displays asymmetric, stem cell-like and oriented cell divisions, which require the activity of the polarly localized protein, BASL. Here we identify the plant-specific BREVIS RADIX (BRX) family as localization and activity partners of BASL. We show that members of the BRX family are polarly localized to peripheral domains in stomatal lineage cells and that their collective activity is required for asymmetric cell divisions. We further demonstrate a mechanism for these behaviors by uncovering mutual, yet unequal dependencies of BASL and the BRX family for each other’s localization and segregation at the periphery of stomatal lineage cells.

## INTRODUCTION

Asymmetric cell divisions (ACDs), which generate daughters of different composition, size and fate, are an important mechanism for generating cell-type diversity and pattern across all kingdoms of life. Differences between daughter cells of a division may result from their differential inheritance of factors prior to cell division, or may be generated through interactions post-division. In plants, there are numerous extrinsic signals that orient or create asymmetries (Fowler and Quatrano, 1997; Geisler et al., 2000; Nakajima et al., 2001 Lukowitz et al., 2004; Cartwright et al., 2009). Evidence for asymmetry generation in mother cells and differential segregation of proteins to their daughters is more limited; however, it is associated with the stem-cell like divisions within the epidermal lineage that leads to the formation of stomata. Stomata are pores in the epidermis of aerial organs that mediate gas exchange between the plant and the atmosphere. In Arabidopsis, the stomatal lineage undergoes reiterative and oriented asymmetric divisions. This phenomenon, coupled with the accessibility of the leaf epidermis, makes the Arabidopsis stomatal lineage a good model system for the study of cell polarity establishment in a multicellular developmental context. In wild-type Arabidopsis, the stomatal lineage is initiated when meristemoid mother cells (MMCs) divide asymmetrically, generating, as their smaller daughter cells, meristemoids. Meristemoids either undergo additional asymmetric divisions to amplify the number of cells in the lineage or differentiate into guard mother cells (GMCs) and ultimately, two guard cells. The larger MMC daughter cell (stomatal lineage ground cell, SLGC), has dual potential: it may expand in size, assume a lobed morphology and differentiate as a pavement cell, or divide asymmetrically to produce secondary meristemoids at later times (Fig. 1A). In the context of epidermal patterning, these asymmetric cell divisions help to prevent the formation of stomata in contact by placing stomatal precursor cells away from each other and by ensuring that sister cells cannot both assume GMC identity (Pillitteri and Dong, 2013).

**Figure 1.**
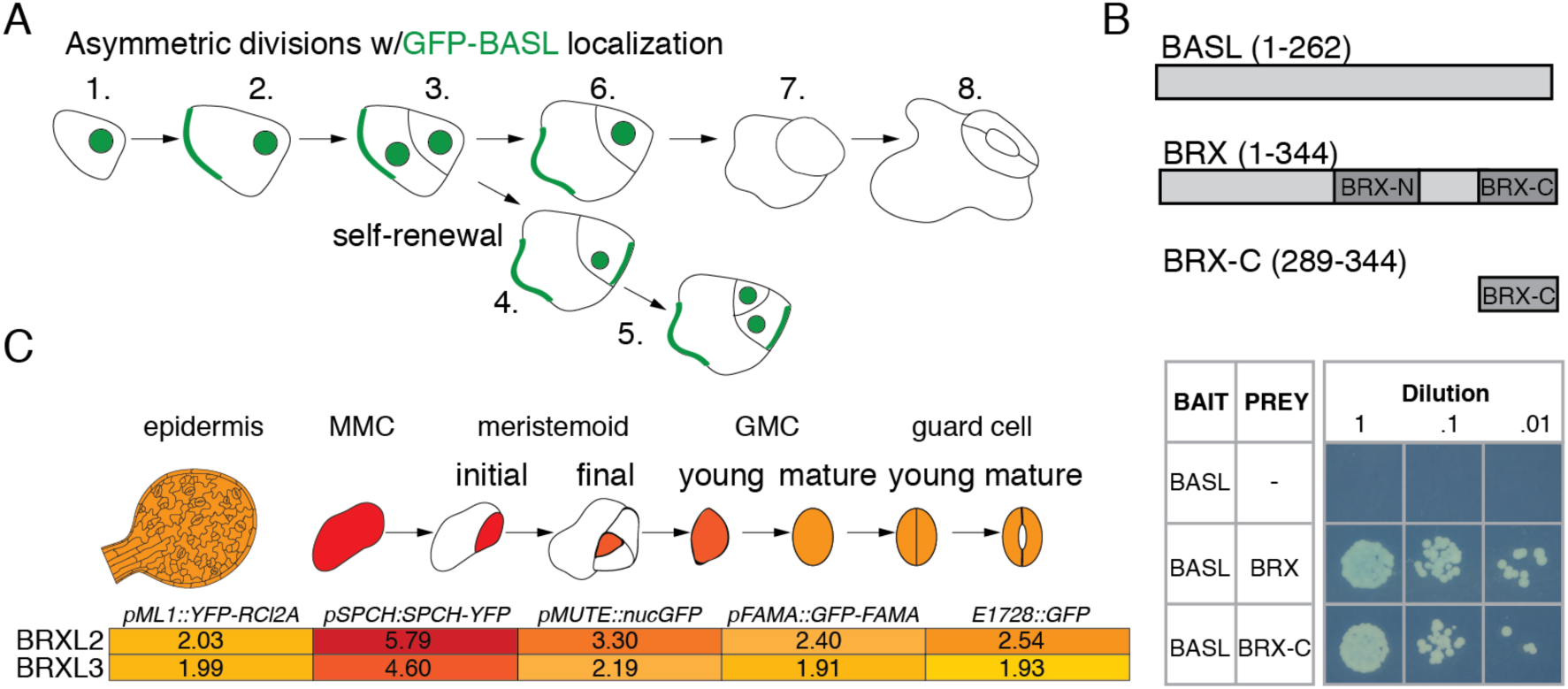
Asymmetric division in the stomatal lineage and identification of BRX family proteins as partners of BASL. **(A)** Scheme of GFP-BASL (green) localization during stomatal lineage divisions. Initially, *BASLp:GFP-BASL* (hereafter, BASL) localizes to nuclei of epidermal cells (1.). As epidermal cells assume MMC identity and expand prior to division, BASL accumulates in a polarized crescent at the cell membrane (2.). Immediately after asymmetric cell division, BASL is expressed in nuclei of both daughters, but peripheral BASL is inherited only by the SLGC (3.). As SLGCs expand in size and become pavement cells, BASL expression fades from the nucleus while its peripheral localization is maintained (4.,6.). BASL peripheral polarization is re-established in secondary MMCs (4.-5.). BASL is no longer present in differentiating GMCs (7.) and stomata (8.). **(B)** Pairwise yeast two-hybrid showing interaction (growth and blue color) between BASL and BRX and BASL with BRX C domain). Numbers in parentheses indicate included amino acid residues. “Bait” and “Prey” denote fusion with GAL4 DNA-binding (BD) and activation (AD) domains, respectively. **(C)** Stage-specific expression enrichment of BRXL2 (At3G14000) and BRXL3 (At1G54180) during stomatal lineage development. Heat maps show unscaled mean and median log2-transformed expression values from Adrian et al., 2015; low expression is in yellow, and high expression is in red.

Regulation of stomatal development involves transcription factors that promote acquisition of and transition through MMC, GMC and guard cell identities and inter- and intra-cellular signaling systems that spatially and temporally limit lineage progression (reviewed in Zoulias et al., 2018). These same components control the expression of factors responsible for executing asymmetric divisions (Lau et al., 2014). A key regulator of asymmetric divisions in stomatal lineage cells is the novel and plant-specific protein BREAKING OF ASYMMETRY IN THE STOMATAL LINEAGE (BASL). BASL polarizes at the cortex prior to division and is differentially inherited by daughter cells in a manner that consistently correlates with the resulting cell identities (Dong et al., 2009). In *basl* mutants, the physical and cell-fate asymmetries of the stomatal ACDs are diminished. The resultant “equalized” sisters hyperproliferate, leading to the accumulation of small stomatal lineage cells and to moderate clustering of stomata (Dong et al., 2009). A second plant specific protein, POLAR LOCALIZATION DURING ASYMMETRIC DIVISION AND REDISTRIBUTION (POLAR), is also differentially segregated in the ACDs in a BASL-dependent manner (Pillitteri et al., 2011). There is evidence that BASL and POLAR act as signaling scaffolds with MAPKs and GSK3β family kinases, respectively. MAPK phosphorylation sites on BASL are needed for BASL to accumulate outside of the nucleus in a polarized cortical crescent, and once activated and polarized, BASL directs the cortical polarization and preferential segregation of MAPK signaling components to one daughter cell (Zhang et al., 2015; Fig. 1A). POLAR affects the nuclear/cytoplasmic ratio of the GSK3β kinase BRASSINOSTEROID INSENSITIVE2 (BIN2), leading to preferential activities at the cell cortex before asymmetric divisions (Houbaert et al., 2018). Both BIN2 and MAPKs target the transcription factor SPEECHLESS (SPCH), leading to its preferential degradation in one daughter of the asymmetric division (Lampard et al., 2008), but BIN2 can also target MAPKs, leading to attenuation of this activity. The balance of kinase distributions, both in terms of cellular and subcellular distribution, is proposed to enable the repeated rounds of asymmetric division and ultimate distribution of differential cell fates in the stomatal lineage.

BASL/MAPK and POLAR/BIN2 contribute to polarity and differential fates, but loss of MAPK activity does not fully depolarize BASL (Zhang et al., 2015) and POLAR appears to act downstream of BASL asymmetries (Pillitteri et al., 2011), indicating that additional components must contribute to stomatal lineage ACDs. Here we identify members of the BREVIS RADIX (BRX) protein family as such components. The family’s founding member, BRX, previously linked to cell proliferation and elongation in the root, has been hypothesized to integrate auxin and brassinosteroid (BR) hormone signaling (Mouchel et al., 2006) and was recently incorporated into a molecular rheostat to modulate auxin flux during protophloem development in the root (Marhava et al., 2018). We demonstrate that in the context of the asymmetric cell divisions of the stomatal lineage, however, BRX and its paralogs are partners of BASL. BRX family members co-localize with BASL in a polar crescent at the cell cortex and are segregated to one daughter; collectively they are required for asymmetric divisions. We further demonstrate a molecular mechanism for these behaviors by showing that BASL and the BRX family share a mutual requirement for each other’s localization and segregation at the periphery of stomatal lineage cells.

## RESULTS

### BRX family proteins are interaction partners of BASL

To identify additional players in BASL-driven stomatal lineage polarity, we integrated a physical interaction screen with stomatal lineage expression surveys. In a yeast two-hybrid screen with full length BASL, ten of 17 positive interactions corresponded to members of the plant-specific BRX family. Subsequent pairwise analyses confirmed the interaction and identified a BRX domain (conserved among all BRX family members) and the central region of BASL (BASL-I) as necessary and/or sufficient for interaction (Figs. 1B and S1A-I). To determine whether this interaction could occur during stomatal lineage divisions, we queried transcriptional profile data and found that *BRLX2* and *BRXL3* transcripts are enriched in stomatal lineage cells (Fig. 1C and Adrian et al., 2015). To confirm the transcriptomic data, we generated transcriptional reporters for each of the five Arabidopsis *BRX* family genes (hereafter, designated *BRXf*) (Fig. S2A-V). *BRXf* gene expression overlaps with *BASL* expression in the epidermis of cotyledons and emerging true leaves (Fig. S2 A-O and Dong et al., 2009) with *BRXL2,* like *BASL,* displaying clear enrichment in asymmetrically-dividing stomatal lineage cells (Fig. S2U-V).

### *BRXf* genes are redundantly required for asymmetric division in the stomatal lineage

*BRX* has a well-established role in root growth (Mouchel et al., 2006) and protophloem differentiation (Rodriguez-Villalon et al., 2014; Marhava et al., 2018). Although a leaf growth phenotype has also been described in *brx* mutants (Beuchat et al., 2010b), this was not investigated at the level of stomatal production or pattern, and no such phenotypes have been reported for other BRXf members in the literature (Mouchel et al., 2004; Briggs et al., 2006). In our own investigation, we found no significant stomatal production or pattern defects in plants bearing the original *BRX*^c^ allele or in single null alleles of *BRXL1, BRXL2,* or *BRXL3* (Fig. S3B-F). We found, however, that multiple mutant combinations exhibit stomatal clusters, hyperproliferation of stomatal lineage cells, and a loss of physical asymmetry among sister cells (Figs. 2A,B and S3G-K). Plants lacking the stomatal lineage-enriched family members *BRXL2* and *BRXL3* exhibit stomatal phenotypes (Fig. S3I, asterisks), but elimination of additional family members as in the *brx*^*C*^ *brxl1 brxl2 brxl3* quadruple mutant (hereafter referred to as *brx-q*) further increases the severity of stomatal defects (Figs. 2B, S3K). Thus, in contrast to the predominant role of the single family member *BRX* in promoting root development (Briggs et al., 2006), there is a clear dosage effect, with multiple *BRX* family members contributing to stomatal development.

**Figure 2.**
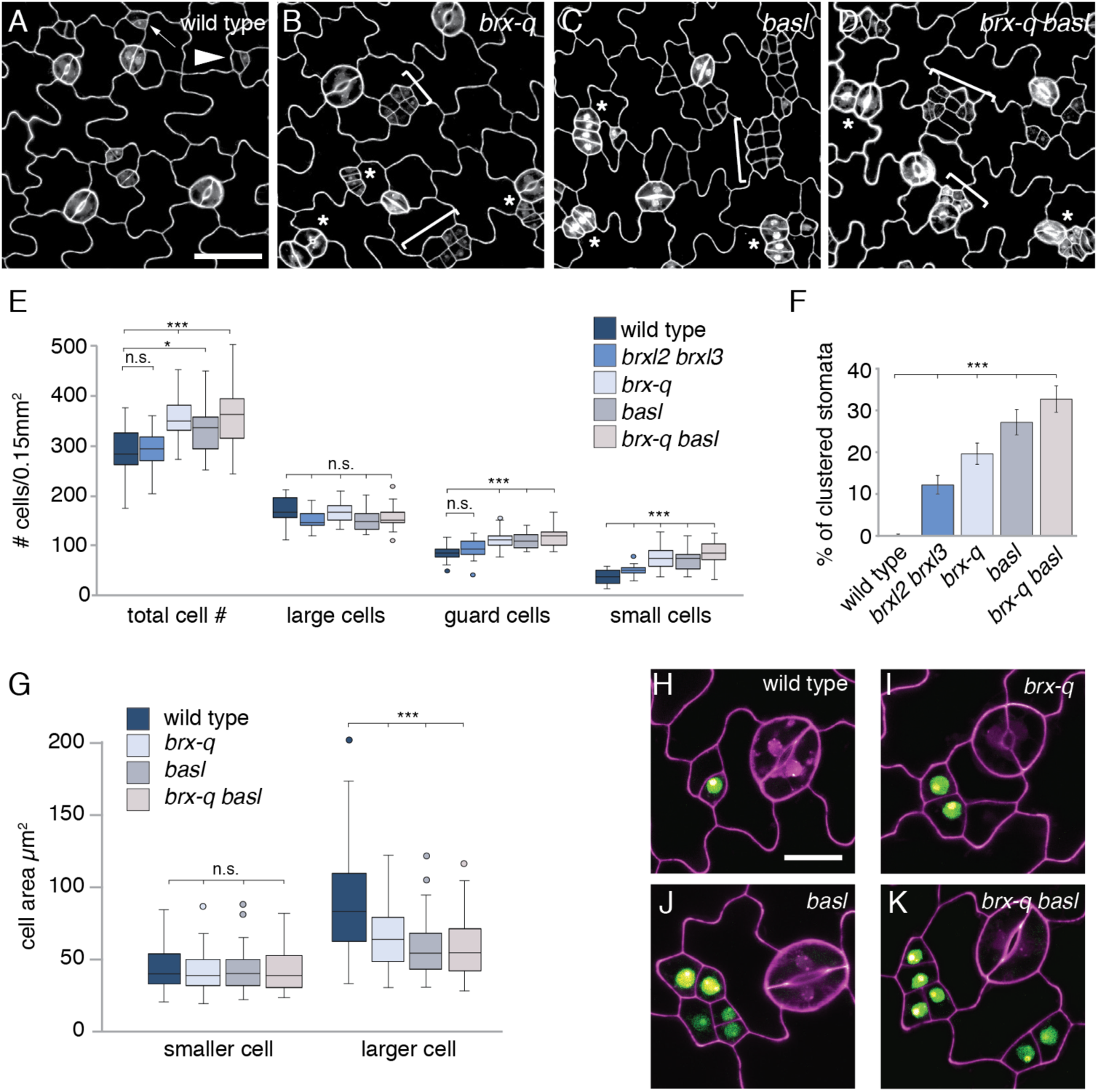
Physical and cell fate asymmetry defects in *brx-q* and *basl* mutants. **(A-D)** Confocal images of 3dpg adaxial cotyledons of wild type, *brx*^*C*^ *brxl1 brxl2 brxl3* (*brx-q*), *basl* and *basl;brx-q*. Cell outlines are visualized by propidium iodide. Arrows and arrowheads indicate meristemoids and SLGCs, respectively. Braces and asterisks indicate clustered small cells and stomata, respectively. Scale bar in A = 50μm; B-D at same scale. **(E)** Quantitation of cell type abundance in 3dpg adaxial cotyledons for the indicated genotypes. For each class, n = 20 plants, *, and *** show significant difference, P < .05, and P < .0001, respectively by two-tailed Wilcoxon rank-sum test for the bracketed comparisons. Center lines in box plots show the medians; box limits indicate the 25th and 75th percentiles; whiskers extend 1.5 times the interquartile range from the 25th and 75th percentiles. **(F)** Quantitation of stomata clustering phenotype in 3dpg adaxial cotyledons for the indicated genotypes. *** shows significant difference, P < .0001, by Fisher’s Exact test for the bracketed comparisons. Error bars indicate the 95% confidence interval. For composition of stomata clusters refer to Fig. S6. **(G)** Quantitation of cell areas for sister-cell pairs of 3dpg adaxial cotyledons. For each class, n = 72 cell pairs. *** show significant difference, P < .001, by two-tailed Wilcoxon rank-sum test for the bracketed comparisons. Center lines in box plots show the medians; box limits indicate the 25th and 75th percentiles; whiskers extend 1.5 times the interquartile range from the 25th and 75th percentiles. **(H-K)** Confocal images of 3dpg adaxial cotyledons expressing *MUTEp:GFP-NLS* (green). Cell outlines were stained with propidium iodide. Genotypes are indicated in panels. Scale bar in H 15μm; I-K at same scale.

### Loss of *BASL or BRXf* leads to similar defects in cell-size and cell-fate asymmetries

The stomatal lineage defects in *brx-q* resembled those in *basl.* We quantified these defects in both mutants compared to WT by measuring pattern defects (clustering) and production of all cell types, binning epidermal cells into three classes: Guard Cells (distinguished by morphology), Large Cells (area > 100 μm^2^), and Small Cells (area < 100 μm^2^). The Large Cells class includes pavement cells and larger SLGCs that are unlikely to divide further while the Small Cells class includes smaller SLGCs, meristemoids, and GMCs. *brx-q, basl*, and *brx-q basl* cotyledons had significantly increased numbers of guard cells and small cells relative to WT (Fig. 2E). Among these mutants, however, there were no significant differences in the number of guard cells and small cells, consistent with *BASL* and *BRXf* genes functioning in a common biological process (Fig. 2B-E). When stomatal clustering is considered, *brxl2 brxl3* mutants significantly differ from wildtype and there is a discernible (though less than additive) difference between *brx-q* and *brx-q basl* (Fig. 2F).

In both *brx-q* and *basl* mutants, size asymmetry between stomatal-lineage sister cells is diminished (Fig. 2B,C). This phenotype could arise from misplacement of the mother cell division plane before division and/or loss of differential cell expansion after division. In the division plane-misplacement scenario, a decrease in the size of one daughter cell would be expected to correlate with an increase in the size of its sister. We measured areas of sister-cell pairs in *basl* and *brx-q* by incorporating the meristemoid identity marker *MUTEp:NLS-GFP* (Dong et al., 2009) to constrain the analysis to meristemoids at a specific developmental stage (Fig. 2G-K). In addition, to focus on cells most recently divided, we used aniline-blue to highlight new walls (Kuwabara and Nagata, 2006; Kuwabara et al., 2011; Fig. S4). With each of these methods, we found that the average area of the larger sister cell was significantly reduced in *brx-q* and *basl* relative to WT, but the average area of the smaller cell was nearly identical across genotypes suggesting that there is a pre-divisional defect in cell size (Fig. 2G, S4B). Given the accumulation of small cells in both *basl* and *brx-q* mutants (Fig. 2B-E), however, additional functions of these factors in post-divisional growth are also possible.

Loss of physical asymmetry corresponds with loss of cell-fate asymmetry between sisters in *brx-q* and *basl* mutants. Mutants accumulate excess small cells, many of which eventually form lobes like pavement cells, but also display excess and clustered stomata (Fig. 2B,C) suggesting that the *BRX* family and *BASL* are required to enforce differential fate decisions rather than to make a specific cell type. This idea is substantiated by gene expression patterns; in WT, expression of *MUTEp:NLS-GFP* is restricted to the smaller sister and marks stomatal precursor identity (Fig. 2H). In *basl, brx-q* and *brx-q basl* mutants, we found that both (or neither) sister cells could express this MUTE reporter (Fig. 2I-K).

### BRX proteins are polarized in stomatal lineage cells

To begin addressing the mechanisms by which *BRXf* proteins function collectively in stomatal development, we generated translational reporters to monitor protein dynamics in the leaf epidermis. We chose members of two distinct family subgroups: BRX, because it is has been most extensively characterized (Beuchat et al., 2010b; Marhava et al., 2018; Mouchel et al., 2006; 2004; Sankar et al., 2011; Scacchi et al., 2009) and BRXL2 because of its stomatal lineage enrichment and non-redundancy with *BRX* in the root context (Beuchat et al., 2010a; Briggs et al., 2006). Like the transcriptional reporter, *BRXL2p:BRXL2-YFP* is detected in MMCs and SLGCs. Remarkably, the reporter localizes to polarized crescents near the plasma membrane (Fig 3A). Polarized crescents of BRXL2-YFP first appear in MMCs and are positioned distal to the division plane such that they are reliably inherited in the larger daughter SLGCs Fig. 3A). BRXL2-YFP polarization is still visible in young expanding SLGCs, but is no longer present in mature pavement cells (Fig. 3A). *BRXp:BRX-YFP* was below the detection limit in leaves, but when expressed under the *BASL* promoter to direct expression in stomatal lineage cells, it polarized similar to BRXL2-YFP (Fig. 3B). Expression of either *BASLp:BRX-YFP* or *BRXL2p:BRXL2-YFP* in a *brx-q* mutant is sufficient to rescue stomatal patterning defects (see Figs. 5B, H), confirming the functionality of the tagged proteins. The polarized peripheral localization of BRXL2-YFP and BRX-YFP strongly resembles that of BASL translational reporters, with the exception of not appearing in nuclei of stomatal lineage cells.

**Figure 3:**
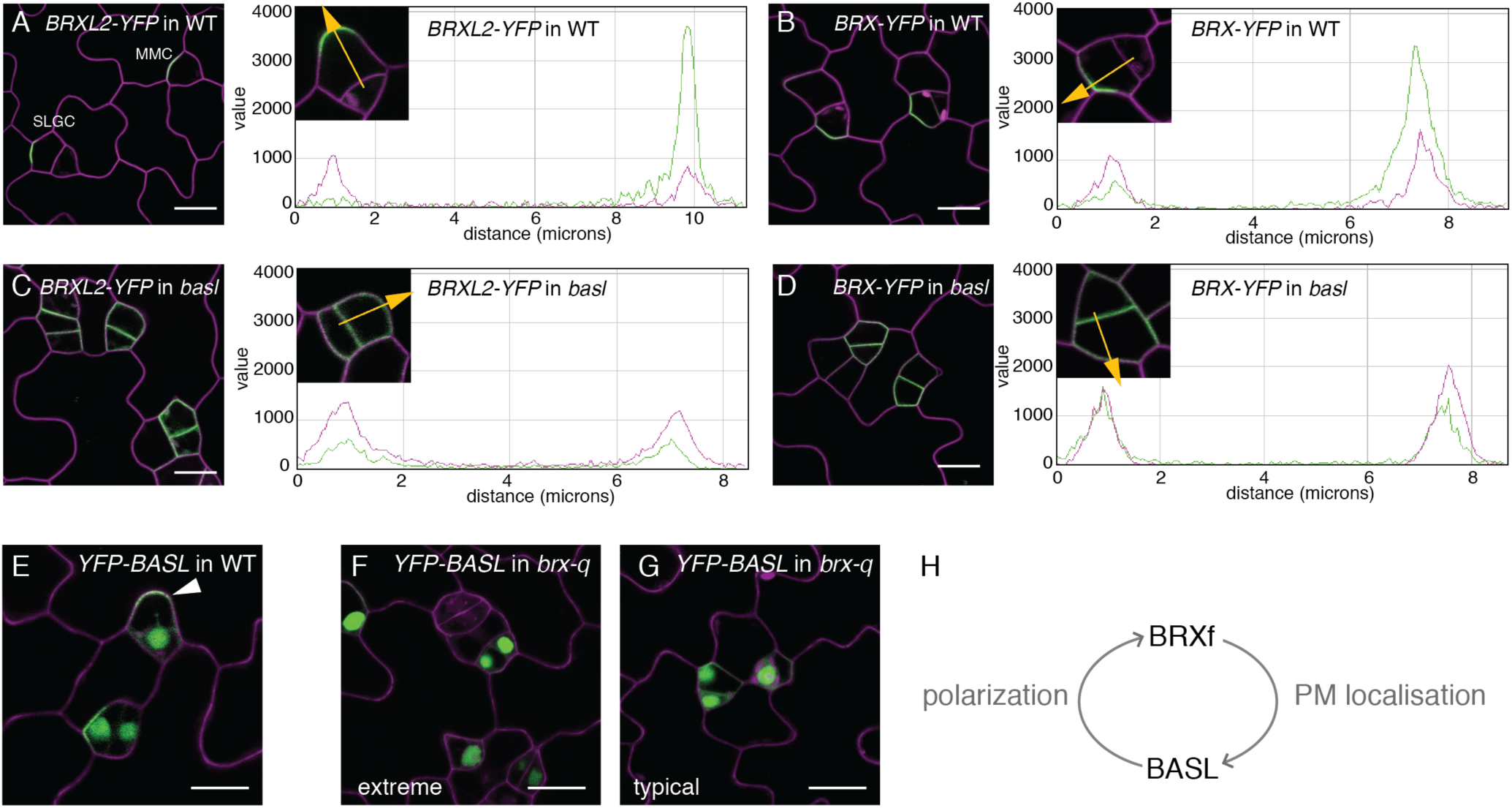
Cortical polarization of BRX family proteins and interdependence of BRXf and BASL for membrane association and polarization. **(A-D)** Confocal images of 3dpg abaxial cotyledons expressing *BRXL2p:BRXL2-YFP* and *BASLp:BRX-YFP* in WT (**A-B**) and *basl* (**C-D**). PIP2A-RFP (magenta) marks the plasma membrane. Quantification of polarity from confocal images of *BRXL2-YFP* and *BRX-YFP* expression (inset) with corresponding plots of pixel intensities along profiles indicated by orange lines. Green and magenta plots represent signal intensities of BRXL2-YFP or BRX-YFP and PIP2A-RFP, respectively. **(E)** *BASLp:YFP-BASL* (green) polarized (white arrowhead) and nuclear in cells of 3-dpg WT abaxial cotyledons (**F-G**) *BASLp:YFP-BASL* exhibiting no (F, 14/95 cells) or diminished (G, 70/95 cells) polarized cortical localization in *brx-q*. Cell outlines (magenta) stained with propidium iodide. Scale bars: 10μm in (A-G). **(H)** Interdependence of BASL and BRXf. BASL is required for BRXf polarization, and BRXf is required for BASL’s localization to the plasma membrane.

**Figure 5.**
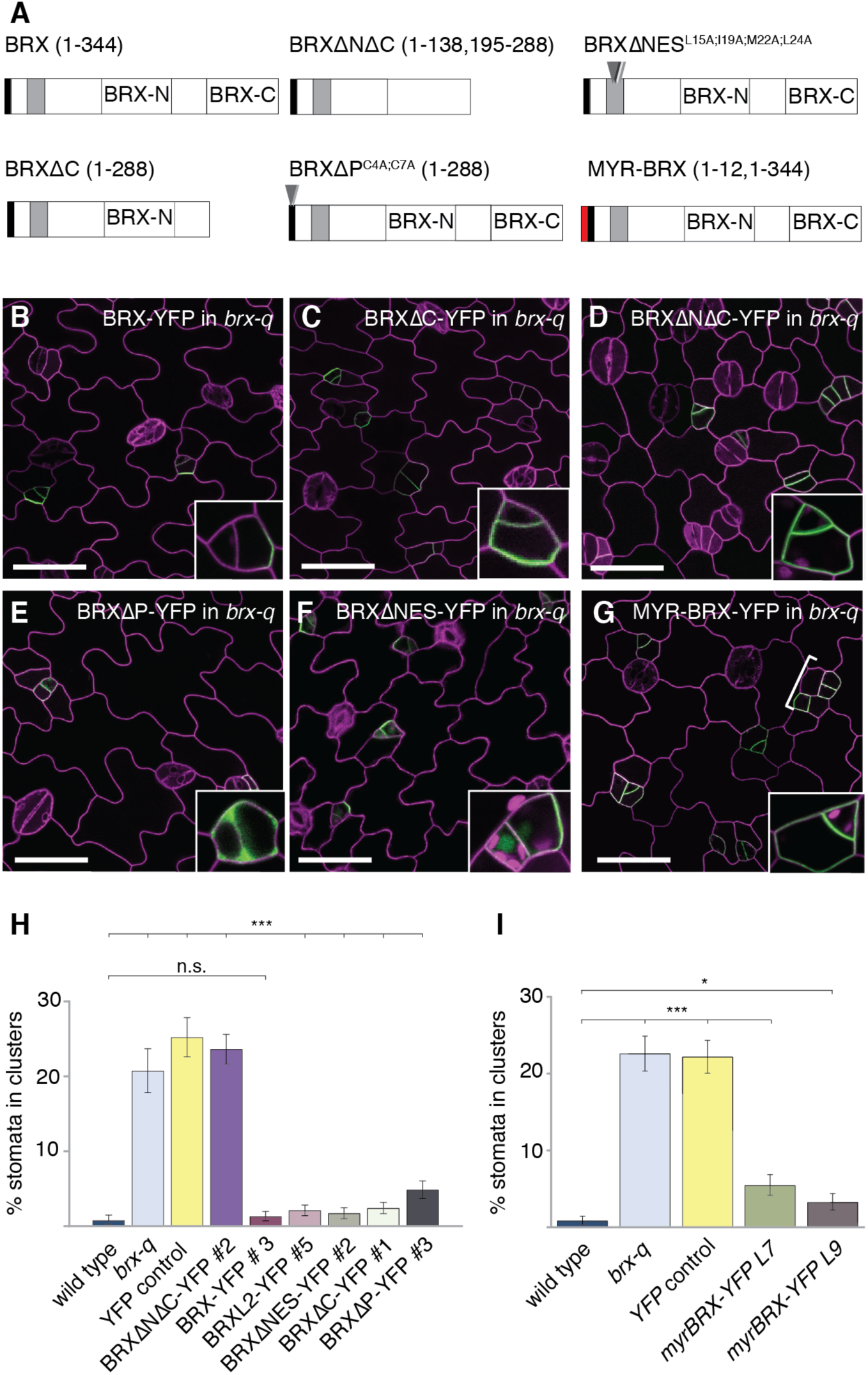
Dissection of protein domains that mediate subcellular localization of function of BRXf during stomatal lineage asymmetric divisions. **(A)** Schematic of modified BRX proteins constructs expressed in B-G. Arrowheads indicate mutated residues and numbers in parentheses are the amino acids included in construct, if not full length (i.e., 344). **(B-G)** Confocal images of 3dpg adaxial cotyledons expressing indicated BRX-variant YFP-tagged reporters. Cell outlines (magenta) visualized with PIP2A-RFP. Insets in each panel show representative protein localization in meristemoid/SLGC pair. (**B**) *BASLp:BRX-YFP* in *brx-q* showing full rescue and polarity. **(C)** *BASLp:BRX*Δ*C-YFP* in wild type. (D) Depolarized and mis-segregated *BASLp:BRX*Δ*N*Δ*C-YFP* in *brx-q* fails to rescue. **(E)** *BASLp:BRX*Δ*P-YFP* in *brx-q* is no longer associated with plasma membrane and its ability to rescue is diminished. **(F)** *BASLp:BRX*Δ*NES-YFP* in *brx-q*. **(G)** *BASLp:myr-BRX-YFP* in *brx-q* **(H)** Quantitation of rescue of stomatal clustering phenotype. For wild type, *brx-q,* and *BASLp:YFP-YFP*; *brx-q*, n = 20. For all other classes, n = 40 and data from 2 independent lines are pooled (n = 20 for each independent line). *** shows significant difference, P < .0001, by Fisher’s Exact test for the bracketed comparisons. Error bars indicate the 95% confidence interval. For composition of stomata clusters refer to Fig. S6. **(I)** Quantification of clustered stomata in wild type and *brx-q*, YFP control (YFP-YFP) in *brx-q*, and myrBRX-YFP in *brx-q*. L7 and L9 are two independent lines. *myr*BRX (partially) rescues the *brx-q* mutant. * and *** shows significant difference, P< .05 and P <.0001, respectively, by Fisher’s Exact test for the bracketed comparisons. Error bars indicate the 95% confidence interval. For composition of stomata clusters refer to Fig. S6. Scale bars in B-G are 50μm.

### BASL and BRX-family proteins are mutually dependent on each other for localization

Physical interaction between BASL and BRXf proteins in yeast and their colocalization in specific stomatal lineage cell-types made us curious about their regulatory hierarchy: could either be responsible for the other’s localization? We found that the polarized cortical localization of BRXL2-YFP or BRX-YFP (Figs 3A,B) was completely lost in *basl* mutants, with BRXL2-YFP and BRX-YFP expanding to mark the entire cell periphery (Figs. 3C,D). In the reciprocal experiment, where we monitored BASL in the absence of BRX family function (*brx-q*), we found instead that the nucleocytoplasmic balance of YFP-BASL was disrupted (Figs. 3E-G). In extreme cases, YFP-BASL localization was strictly nuclear (14/95 cells) (Fig. 3F), but typically, faint cytoplasmic expression and traces of polarized accumulation were also observed at the membrane of dividing MMCs and prospective SLGCs (70/95 cells) (Fig. 3G) and occasionally BASL appeared normally polarized (11/95 cells). These observations indicate that the BRXf and BASL participate in an unequal mutualistic relationship in which the BRXf promotes accumulation of the proteins at the plasma membrane and BASL is responsible for polarization at this site (Fig. 3H).

### BRX and BASL interact physically and affect one another’s localization *in vivo*

To monitor BRX and BASL behavior simultaneously, we crossed BRX-CFP and BASL-YFP lines. In the F2 progeny of this cross, plants bearing a single reporter behaved as observed previously, with BRX-CFP polarized at the membrane in MMCs and SLGCs (Fig. 4A) and BASL-YFP exhibiting nuclear and polarized localization in the same cell types (Fig. 4B). In plants expressing BRX-CFP and BASL-YFP simultaneously, the proteins completely colocalize at the cell periphery (Figs. 4C-E) with mean values >0.9 for Pearson’s and Manders’ coefficients (Fig. 4P) (Manders et al., 1992). In plants expressing both proteins, we also observed that BRX-CFP polarizes to a greater extent than it does in the absence of exogenous BASL (Figs. 4A,C). Unexpectedly, when in a WT background, the presence of BRX-CFP prevents BASL-YFP from accumulating in nuclei (Figs. 4B, 4D). This appears to be due to excessive BRX because when the same reporters are monitored in *brx-q* mutant, BASL-YFP can be seen in the nucleus (Fig. 4G-H). Taken together, the peripheral polarization of BRX-CFP and the nuclear accumulation of BASL-YFP appear to be titrated by levels of the other protein.

**Figure 4.**
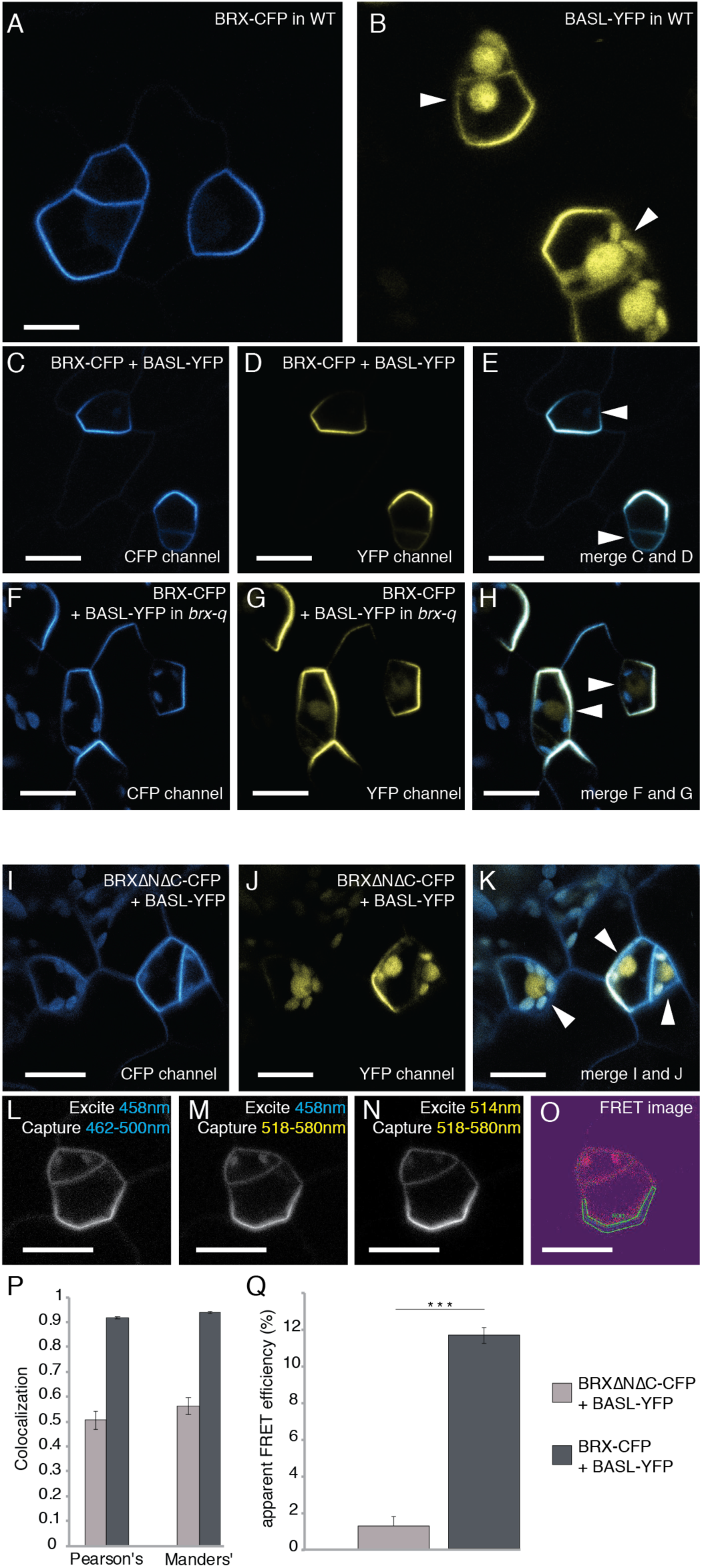
Demonstration of BRX and BASL complex formation *in vivo* (A-E) Confocal images of wild-type abaxial cotyledons at 3dpg expressing BASL promoter driven BRX-CFP variants (cyan) and/or BASL-YFP (yellow). **(A)** BRX-CFP alone **(B)** BASL-YFP alone. Arrowheads indicate BASL-YFP nuclear localization. **(C)** BRX-CFP localization in the presence of BASL-YFP. **(D)** BASL-YFP localization in the presence of BRX-CFP. **(E)** Merge of (C) and (D). Arrowheads indicate absence of nuclear BASL-YFP. **(F-H)** 3-dpg *brx-q* abaxial cotyledons expressing *BASLp:BRX-CFP, BASL-YFP* and merge, arrowheads in H indicate weak BASL-YFP in nuclei. **(I-K)** Colocalization of BRXΔNΔC-CFP, BASL-YFP and merge arrowheads in (**K**) indicate strong BASL-YFP nuclear localization in the presence of BRXΔNΔC-CFP. **(L-O)** FRET between *BASLp:BRX-CFP* (FRET donor) and BASL-YFP (FRET acceptor). Typical series of images generated for measurement of FRET by the sensitized emission method, with excitation in emission capture spectra displayed on image. **(O)** Corrected FRET image obtained by pixel-by-pixel calculation of FRET intensities from images (L-N). Correction factors were obtained by imaging cells expressing BRX-CFP or BASL-YFP in isolation. Intensity values range from 0 to 1 and are indicated by a color spectrum of purple to red. Pixels within the indicated ROI were included in the calculation of apparent FRET efficiency for this particular sample (details in methods). **(P)** Quantitation of Pearson’s and Manders’ colocalization coefficients measured from images of cells expressing combinations of BRX-CFP or BRXΔNΔC-CFP with BASL-YFP. For each class n=15 cells. Error bars represent standard error of the mean. **(Q)** Quantitation of apparent FRET efficiencies measured from images of cells expressing combinations of BRX-CFP or BRXΔNΔC-CFP with BASL-YFP. For each class n=50 cells. Error bars represent standard error of the mean. *** signifies P < .001 by two-tailed Wilcoxon rank-sum test for the bracketed comparisons. Scale bars: 10μm in (A-O).

Functional and localization data suggest that BASL and BRX form a complex; to test complex formation in the stomatal lineage, we wished to use Förster Resonance Energy Transfer (FRET) to test protein interactions *in vivo*. To interpret FRET data, however, we needed to create a non-interacting control that was still present in the same compartment of the cell. Based on the Y2H data that indicated the BRX-domains as the sites of interaction, we created a BRX protein missing both BRX domains (BRXΔNΔC). This yielded a protein that exhibited peripheral localization in stomatal lineage cells (Fig. 4I) but could not dominantly inhibit BASL-YFP nuclear accumulation (Figs. 4J,K, 4P). FRET analyses were then performed on stomatal lineage cells co-expressing both BRX-CFP and BASL-YFP (and BRXΔNΔC-CFP with BASL-YFP) through the measurement of sensitized emission (van Rheenen et al., 2004; Figs. 4L-O). BRX-CFP and BASL-YFP exhibited an average apparent FRET efficiency of 11.7% (Fig. 4Q). This is significantly different from the FRET efficiency (1.3%, P=<.001) observed between BASL-YFP and BRXΔNΔC-CFP (Fig. 4Q). Thus, in stomatal lineage cells, BRX-CFP and BASL-YFP are likely part of the same protein complex.

### BRXf localization and function requires BRX domains and plasma membrane association via a palmitoylation motif

Similar to stomatal lineage-expressed and polarized proteins BASL and POLAR, BRXf proteins lack domains of clearly defined biochemical function, but do contain sites that mediate protein interactions. Deletion of either the N-terminal (BRXΔN) or the C-terminal (BRXΔC) BRX domain alone reduced, but did not completely abrogate the protein’s ability to polarize, influence BASL localization, or rescue stomatal patterning defects in *brx-q* mutants (Figs. 5A,C and Fig. S5C,H). In contrast, deletion of both BRX domains (BRXΔNΔC) yielded a protein that completely failed to polarize or rescue stomatal clustering in *brx-q* (Fig. 5A,D and Fig. S5D). Interestingly, even the BRXΔNΔC protein still appeared associated with the plasma membrane, suggesting that other domains mediate subcellular localization (Fig. 5D). BRXf proteins have putative palmitoylation signals at their N-termini, (Fig. S5A, orange box). Palmitoylation is a reversible lipid modification associated with membrane localization of polarized proteins such as human R-RAS (Baumgart et al., 2010) and *Arabidopsis* Rho of Plants (ROPs) (Sorek et al., 2010). Mutation of the two cysteines in the putative palmitoylation site of BRX (C^4,7^→A^4,7^ or BRXΔP, Fig. 5E) severely reduced membrane localization and polarization (Fig. 5E) and significantly compromised rescue of *brx-q* phenotypes (Figs. 5H and Fig. S5G,H), suggesting that palmitoylation is important for proper localization and function of BRXf proteins.

Although BRX has been proposed to have a nuclear function in protophloem and can be localized there under auxin treatments (Scacchi et al., 2009; Marhava et al., 2018), we did not detect BRX nor BRXL2 in the nucleus in the stomatal lineage under normal conditions (e.g. Fig. 3A-D). We did notice, however, that BRXf proteins encode a predicted NES. To determine whether there was a role for BRXf in the nucleus in the stomatal lineage, we removed the NES and assayed localization and function and, conversely, we prevented BRX from entering the nucleus by tethering to the plasma membrane using an N-terminal myristoylation signal (Roy et al., 2002). BRXΔNES-YFP driven by the BASL promoter can accumulate in the nucleus of stomatal lineage cells (Fig. 5F), but was indistinguishable from full length BRX in its capacity to rescue the *brx-q* mutant (Fig. 5H). Expression of BRXΔNES-YFP also did not create any obvious gain of function phenotype in the stomatal lineage or in leaf growth (not shown). The Myr-BRX-YFP construct exhibited tight plasma membrane localization (Fig. 5G), and was also able to rescue *brx-q* phenotypes (Fig. 5I), suggesting that it is the polarized peripheral localization of the BRX family that is functional during the asymmetric divisions in the stomatal lineage.

### Evolution of the BRX-BASL interaction

We have shown that in the Arabidopsis stomatal lineage, BASL and BRXf proteins are highly interdependent; they require each other for polarized peripheral localization and mutants exhibit similar defects in cell size and fate asymmetry. This dependency of BRXf on BASL activity is particularly intriguing because of the evolutionary history of the proteins. Whereas *BRX* family members can be identified in numerous plant lineages including other dicots, grasses and the moss Physcomitrella (Briggs et al., 2006), *BASL* is only reliably identified in the Brassicacae (Phytozome v.12). This suggests that a BASL-BRXf complex is a recent invention and that BASL’s appearance added a new functionality to BRX, recruiting it from other complexes and activities. Because BRX is normally expressed in a polarized fashion in root protophloem, a pattern linked to PIN-mediated auxin transport (Marhava et al., 2018), we tested whether this (presumably older and more widespread) activity persists in the absence of *BASL*. Such was the case – *BRXp:BRX-YFP* was equally polarized in WT and *basl* mutant roots (Fig. 6A-B). We then addressed whether the information sufficient to enable polar peripheral localization is intrinsic to these two proteins by attempting to reconstitute BRX and BASL polarization in a heterologous system (Fig. 6C). We exploited *S. cerevisiae* as a naïve host and expressed YFP-BASL under a constitutive promoter, then induced BRX expression and monitored YFP dynamics for at least two cell cycles. YFP-BASL initially localizes to the yeast nucleus, but upon induction of BRX, YFP-BASL accumulates at the cell periphery (Fig. 6C). Despite this marked shift in YFP-BASL localization, we did not observe polarization of the proteins, indicating that other, potentially plant-specific, factors are required.

**Figure 6.**
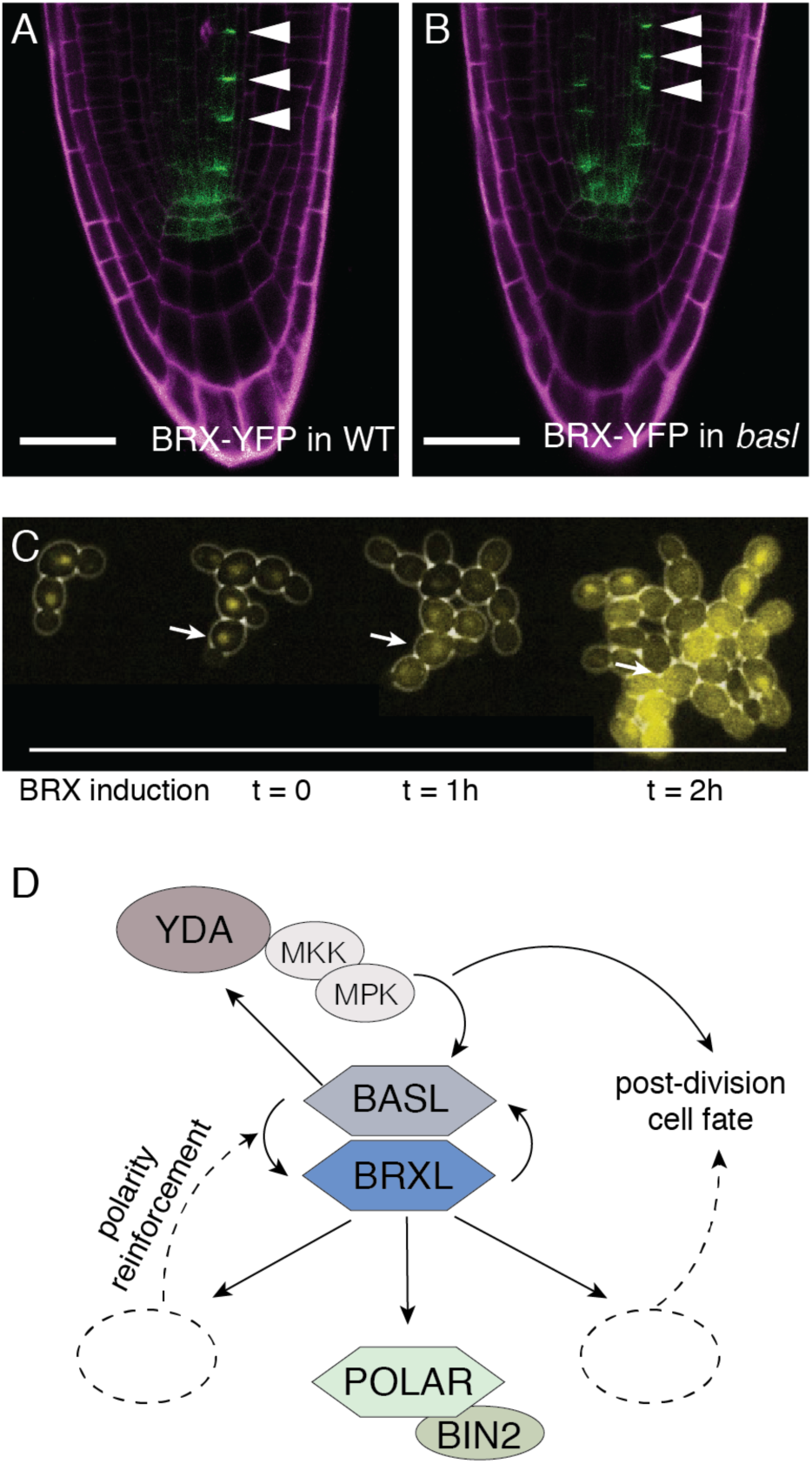
Evidence for and model of novel recruitment of BRXf activities by BASL in the context of Arabidopsis stomatal lineage development. (A-B) Confocal images of 3dpg roots expressing *BRXp:BRX-YFP* (green), cell outlines visualized with propidium iodide (magenta). Arrowheads indicate BRX-YFP polarization in vascular cells in both WT (A) and in *basl* (B) Scale bar 30μm. (C) Reconstitution of BRX-BASL interactions and subcellular localization in yeast. (D) Model for BASL and BRXf interactions in polarity establishment in *Arabidopsis* stomatal lineage cells.

## DISCUSSION

Arabidopsis stomatal lineage cells are among many diverse organs and organisms that generate cell polarity in connection with formative and oriented asymmetric cell divisions. In contrast to other examples found in plants, neither stomatal lineage divisions, nor the polarity proteins marking them e.g., (Yoshida et al., 2019), are absolutely aligned with major body axes (Bringmann and Bergmann, 2017; Mansfield et al., 2018). In contrast to animals and yeast, stomatal lineage cell polarity involves novel proteins. In this work, we identified members of the BRX family as partners for the Arabidopsis stomatal lineage polarity factor BASL. BASL and BRXf establish cell size and fate divergence between daughter cells. Qualitative and quantitative similarities between the *basl* and *brx-q* mutant phenotypes and the lack of additivity observed in *brx-q basl* quintuple mutants indicate that their roles in asymmetric cell division involve a common biological process, an idea further supported by BASL and BRXf co-localization in polarized peripheral crescents. The mutual, but unequal dependence of BASL and the BRXf for each other’s subcellular and polarized localization and previously established physical and genetic relationships between each protein and others suggest that BRX and BASL are part of a core polarity generating system. This system enables the scaffolding, positioning and segregation of other proteins with important functions in creating cell size asymmetries pre-division and executing or amplifying cell fate differences post division (Model in Figure 6D).

Scaffolding is an emerging theme in the stomatal lineage, with two plant-specific proteins with no discernable biochemical activity, BASL and POLAR, each associating with a powerful and broadly used intracellular signaling cascade to provide cell-type and asymmetric division specific behaviors. BASL depends on MAPK phosphorylation for its peripheral polarization before asymmetric division, then, biases the capacity for MAPK signaling towards the SLGC post-division by sequestering the upstream MAPKKK, YODA in the larger daughter (Zhang et al., 2015). POLAR, through its binding of BIN2, changes the nuclear/cytoplasmic balance of the latter, with consequences to MAPK signaling and ultimately to the stability of SPCH in meristemoids post-division (Houbaert et al., 2018). Both MAPK and BIN2-related signaling cascades are used in a large number of developmental and physiological events, and other scaffolds have been identified, perhaps most interesting among them is OCTOPUS (OPS) that, similar to POLAR, sequesters BIN2 away from the nucleus from its a polarized plasma-membrane location in protophloem cells (Vatén et al., 2018). A question that remains, however, is whether polar cortical localization is only a means to ensuring segregation during division (BASL/MAPK) or sequestration from the nucleus (POLAR or OPS/BIN2), or whether polarization there are activities that must take place in a restricted region of the cortex of developing cells.

The BRX family, too is only found in plants and encodes proteins whose sequence and structure give little hint to their activity. Could this family also act as scaffolds, or co-scaffolds with BASL? If so, who are the clients? Extensive studies over the last decade showed that BRX promotes the accurate and coordinated timing of cell fate commitment in the protophloem sieve elements of the root vasculature (Scacchi et al., 2009; Depuydt et al., 2013; Rodriguez-Villalon et al., 2014; Marhava et al., 2018). Although initially reported as a transcriptional regulator (Mouchel et al., 2006), and reported to accumulate in the nucleus under auxin treatment (Briggs et al., 2006; Scacchi et al., 2009), BRX has also been found polarly distributed in protophloem cells, coincident with PIN auxin transporters (Scacchi et al., 2009; 2010). With these two potential activities in mind, our experimental dissection of BRX protein domains and tethering BRX to the plasma membrane firmly suggest that in the stomatal lineage, it is the polarized peripheral BRX pool that is critical for regulating cell fate and divisions. Our identification of the function for BRX’s C-terminal palmitoylation motif is also interesting in light of recently work proposing an “auxin efflux rheostat” model involving BRX and the kinase PROTEIN KINASE ASSOCIATED WITH BRX (PAX) (Marhava et al., 2018). A critical feature of the rheostat model is for BRX to be reversibility associated with the plasma membrane. Of the common lipid modification on proteins, only palmitoylation (S-acylation) is reversible, thus creating a rationale for this specific mode of membrane tethering.

With multiple potential partners in different parts of the cell, a generally unstructured protein sequence interrupted by two BRX domains capable of protein-protein interactions, BRXf proteins do have hallmarks of classical scaffolds. Does the BRX family work with BASL in regulating MAPKs, or does it scaffold yet another kinase family in the stomatal lineage? There is currently no evidence for direct BIN2-BRXf interactions (Houbaert et al., 2018), nor for MAPK-BRXf interactions (J. Dong, pers. comm.), and is interesting that the region of BASL that interacts with BRX (BASL-I, Figure S1E) is independent of its MAPK docking domain and polar membrane association domain (Zhang et al., 2015; 2016).

Whether there is a stomatal PAX-like protein has yet to be shown, but there also are important cautions in analogizing BRX behavior in protophloem with that in the stomatal lineage. *BRX* itself is only weakly expressed in the stomatal lineage and its role in asymmetric division and cell fate is only revealed in multiple mutant combinations that include the stomatal lineage enriched *BRXL2* and *BRXL3* genes. Expression of *BRXL2p:BRXL2-YFP* is sufficient to rescue the *brx-q* mutant in terms of stomatal development, but overexpression of *BRXL2* cannot substitute for lack of *BRX* in the root (Briggs et al., 2006). Additionally, while BRX and BRXL2 are dependent on *BASL* for polar localization in the stomatal lineage, this is not true in the root vasculature. Thus, biochemically and developmentally, BRX appears to do something that BRXL2 cannot in one tissue, and the relationship with BASL in the stomatal lineage may provide some unique functionalities in another.

## METHODS

### Plant Materials

Throughout this study, the *Arabidopsis thaliana* ecotype Columbia-0 was used as a wild-type. Previously published lines used in this study are: *brx*^*C*^, *brxl1, brxl2, brxl3* (Briggs et al., 2006; Mouchel et al., 2004), *basl*-2, pMUTE::NLS-GFP, pBASL::GFP-BASL (Dong et al., 2009) and p35S::PIP2A-RFP (Nelson et al., 2007). *brx*^*C*^ *brxl1 brxl2* and *brx*^*C*^ *brxl3* multiple mutants (gifts from Christian Hardtke) were crossed together to generate the *brx*^*C*^ *brxl1 brxl2 brxl3* quadruple mutant (*brx-q*), which was subsequently crossed with *basl-2* to generate the *brx-q basl* quintuple mutant. AGI codes: BASL (At5g60880), BRX (At1g31880), BRXL1 (At2g35600), BRXL2 (At3g14000), BRXL3 (At1g54180), BRXL4 (At5g20540). Genotypes were confirmed by PCR-based genotyping (primers in Fig. S7). To permit the most accurate comparison of expression patterns, well-characterized reporters were incorporated into the various genetic backgrounds by introgression rather than transformation. In the case of *brx-q* and *brx-q basl* multiple mutants, this approach was not feasible and instead, reporters were transformed in and at least two independent transformed lines scored for expression or phenotypes. Seedlings were germinated on half-strength MS agar plates in a Percival incubator with 16 hr light/8 hr dark cycles for 7-10 days at 22°C. Plants were transferred to soil for growth in a 22°C growth room with 16 hr light/8 hr dark cycles.

### DNA Manipulations

The majority of DNA manipulations involved implementation of vectors compatible with Gateway cloning technology (Invitrogen). GUS transcriptional reporters were generated by cloning 5’ regulatory regions (amplified from genomic DNA) into pENTR D-TOPO (Invitrogen), followed by transfer into the plant binary vector pMDC163 (Curtis and Grossniklaus, 2003). BRX-family translational fusions were generated by cloning corresponding cDNAs into pENTR. Template for BRX cDNA was ABRC stock G20350 and template for BRXL2 was 7dpg seedling WT cDNA. BRXL2 and BASL promoters (from genomic DNA) were cloned into pENTR NotI site before recombination into pHGY (Kubo et al., 2005). BRX functional domains mutants were generated by PCR of BRX pENTR using mismatch or fusion primers. BASLpYFP-BASL was generated by double recombination of the BASL promoter and a YFP-BASL fragment into pGWB510 (Nakagawa et al., 2008). Transgenes were integrated into plants via Agrobacterium-mediated transformation (strain GV3101) according to standard protocols and transformants were selected on the basis of antibiotic resistance and/or observed transgene expression.

### Yeast two-hybrid screen and tests

A yeast two hybrid screen was performed using the full length BASL as bait and a library derived from seedling cDNA (ABRC stock CD4-22). Of the 17 positive colonies identified, 2, 7, and 1 clone(s) of BRXL2, BRXL3 and BRXL4, respectively, were identified. Directed yeast two-hybrid constructs were generated by ligation of cDNAs into vector components of the Matchmaker II system (Clontech). Bait and prey clones were cotransformed into yeast strain AH109, and physical interactions were assayed according to the manufacturer’s protocols.

### Phenotypic Analysis

Scoring of clustering and production of small cells was performed as described in Dong et al., 2009. Scoring of epidermal patterning phenotypes was performed on confocal images of 3dpg adaxial cotyledons. For each genotype, images of .1mm^2^ regions were acquired from the same quadrant of 20 cotyledons and scored blind (scorer did not know the genotype). Cells were binned into three categories on the basis of size (small cells < 100μm^2^ and large cells > 100μm^2^) and morphology (guard cells). Cell surface areas were measured using the “freehand selection tool” and the “measure” function in the ImageJ software package (National Institutes of Health; NIH). Statistical analyses were performed in Excel (Microsoft). Imaging and measurement of sister cell pairs (n = 72 for each genotype) expressing the *pMUTE::NLS-GFP* reporter were similarly performed.

Rescue of stomatal clustering was scored on DIC images of 7dpg abaxial cotyledons cleared in a 7:1 (v:v) EtOH:Acetic Acid solution and mounted in Hoyer’s medium. For each transgenic line, images of .320mm^2^ regions were acquired from the same quadrant of 20 cotyledons and scored blind. At least 2 independent transgenic lines were included for each construct and a *pBASL::YFP-YFP* transcriptional reporter was used as a negative control.

### Microscopy, Image Processing and Analysis

A Leica DM2500 microscope and DFC425 camera were used to obtain DIC images. Confocal images were generated on a Leica SP5 CLSM using sequential line scanning. All image processing was performed in ImageJ (National Institutes of Health; NIH). Colocalization analysis was performed on background-subtracted images using the “Manders’ Coefficients” ImageJ plugin (Manders et al., 1992; Schneider et al., 2012). Profile plots of pixel intensities were obtained on background-subtracted images using the “multi-plot” function in the ImageJ ROI manager. FRET analysis was performed using the “FRET SE Wizard” application included in the Leica Application Suite software package. CFP-tagged donor molecules were excited by a 458nm laser and donor emission was gated at a spectral range of 462-500 nm. YFP-tagged acceptor molecules were excited by a 514nm laser and acceptor emission (for both direct excitation and FRET) was gated at a spectral range of 518-580 nm. Images of cells expressing BRX/BRXΔNΔC-CFP (donor only) or BASL-YFP (acceptor only) in isolation were obtained to generate correction factors for donor cross-talk in the acceptor channel and acceptor cross-excitation by the 458nm laser. Apparent FRET efficiency was calculated according to the equation: (B-Aβ - C(γ - αβ))/C(1 - βδ) (van Rheenen et al., 2004) where A, B, and C are signal intensities for the Donor, FRET, and Acceptor channels, respectively, β is a correction factor obtained from the donor-only image, and α, γ, and δ are correction factors obtained from the acceptor-only image.

## SUPPLEMENTAL INFORMATION

Figure S1. Pairwise yeast two-hybrid analysis of BRX/BASL

Figure S2. Transcriptional reporters of *BRX* family

Figure S3. Additional stomatal phenotypes of *BRX* mutants

Figure.S4. Aniline-blue analysis of cell size asymmetries

Figure S5. BRX sequence modifications affect its localization

Figure S6. Details of stomatal clustering in Figures 2 and 5

FigureS7. Description of primers used

### ACKNOWLEDGMENTS

We thank Christian Hardtke (University of Lausanne) for discussions and *BRX* alleles, Tsuyoshi Nakagawa for transformation vectors, David Ehrhardt (Carnegie, DPB) for microscopy advice and members of our research group for numerous discussions and comments on the manuscript. Initial work was supported by NSF grant IOS-084552. M.R. was partially supported by NIH training grant 5 T32 GM007276 to Stanford University. D.B is an investigator of the Howard Hughes Medical Institute

## AUTHOR CONTRIBUTIONS

M.R., J.D. and D.B. designed wet-lab experiments and analyzed results. M.R. and J.D. performed experiments. M.R., A.W. and D.B. wrote the manuscript.

