## Supplemental Figure 1-7 for "A Plant-Specific Polarity Module Establishes Cell Fate Asymmetry in the Arabidopsis Stomatal Lineage"

### SUPPLEMENTAL FIGURES 1-7

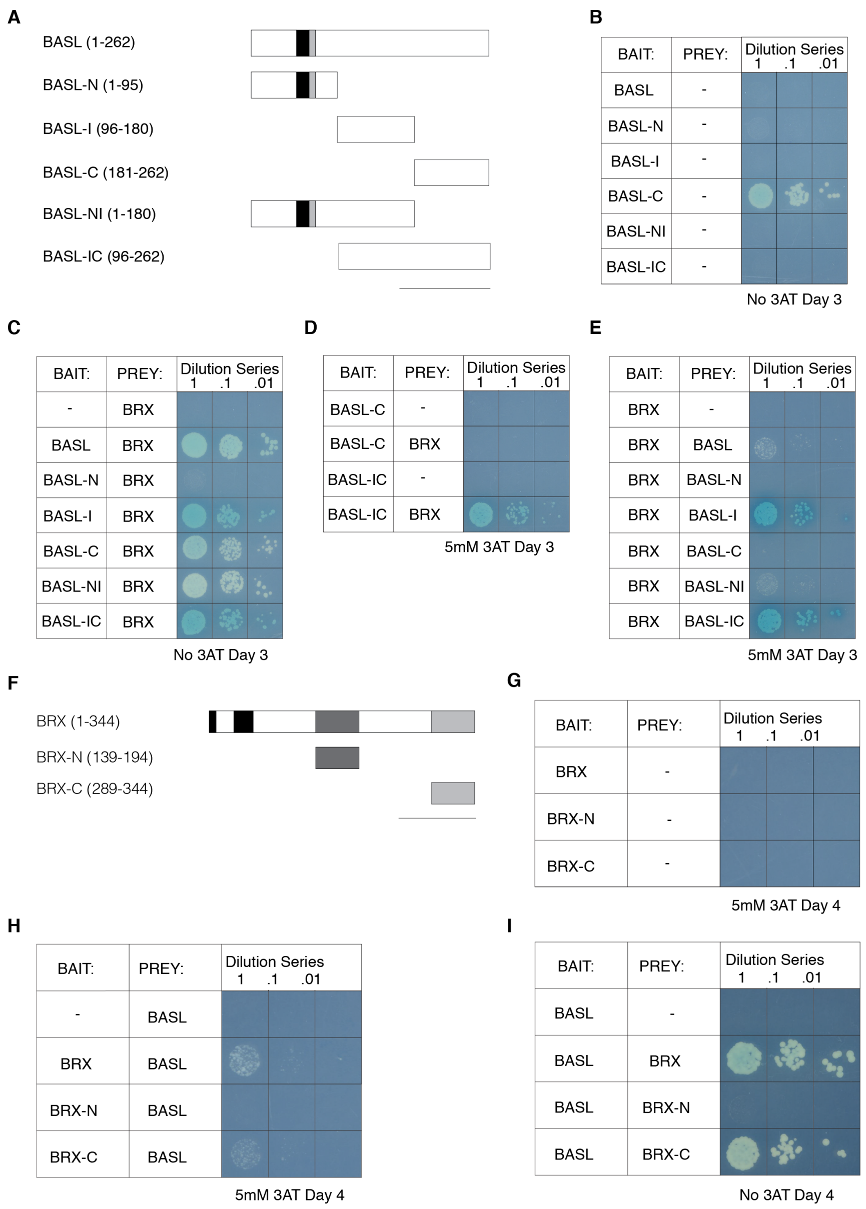


**Figure S1. Pairwise yeast two-hybrid analysis of physical interaction between BRX and BASL (A)** Schematic of full-length BASL peptide and truncations used in yeast two-hybrid analyses. Predicted nuclear localization (NLS) and export (NES) signals are denoted in black and gray, respectively. The amino acids included in the construct are in parentheses. Scale bar is equivalent to 100 amino acids. **(B-E)** Pairwise yeast two-hybrid analysis of physical interaction between full-length BRX and BASL (and BASL truncations). In this analysis "bait" and "prey" were fused to GAL4 activation (AD) and DNA-binding (BD) domains, respectively. Yeast were grown for 3 days on –His minimal media containing X-α-Gal. 3AT was included in concentrations indicated below panels. **(B)** Analysis of BD-BASL autoactivity in yeast carrying an empty version of the AD-vector. **(C)** Analysis of physical interaction between BD-BASL and AD-BRX protein fusions. **(D)** Inhibition of BD-BASL-C autoactivity with 5mM 3AT. (E) Analysis of physical interaction between BD-BRX and AD-BASL protein fusions (reciprocal of C). **(F)** Schematic of full-length BRX peptide and truncations used in yeast two-hybrid analyses. Domains conserved among BRX family genes in Arabidopsis are denoted in black, two BRX domains in grey. Scale bar is equivalent to 100 amino acids. **(G-I)** Pairwise yeast two-hybrid analysis of physical interaction between full-length BASL and BRX (and BRX domains). Yeast were incubated for 4 days, and in the presence of 3AT where indicated. **(G)** Inhibition of BD-BRX autoactivity with 5mM 3AT. **(H)** Analysis of physical interaction between BD-BRX and AD-BASL protein fusions. **(I)** Analysis of physical interaction between BD-BASL and AD-BRX protein fusions (reciprocal of H).

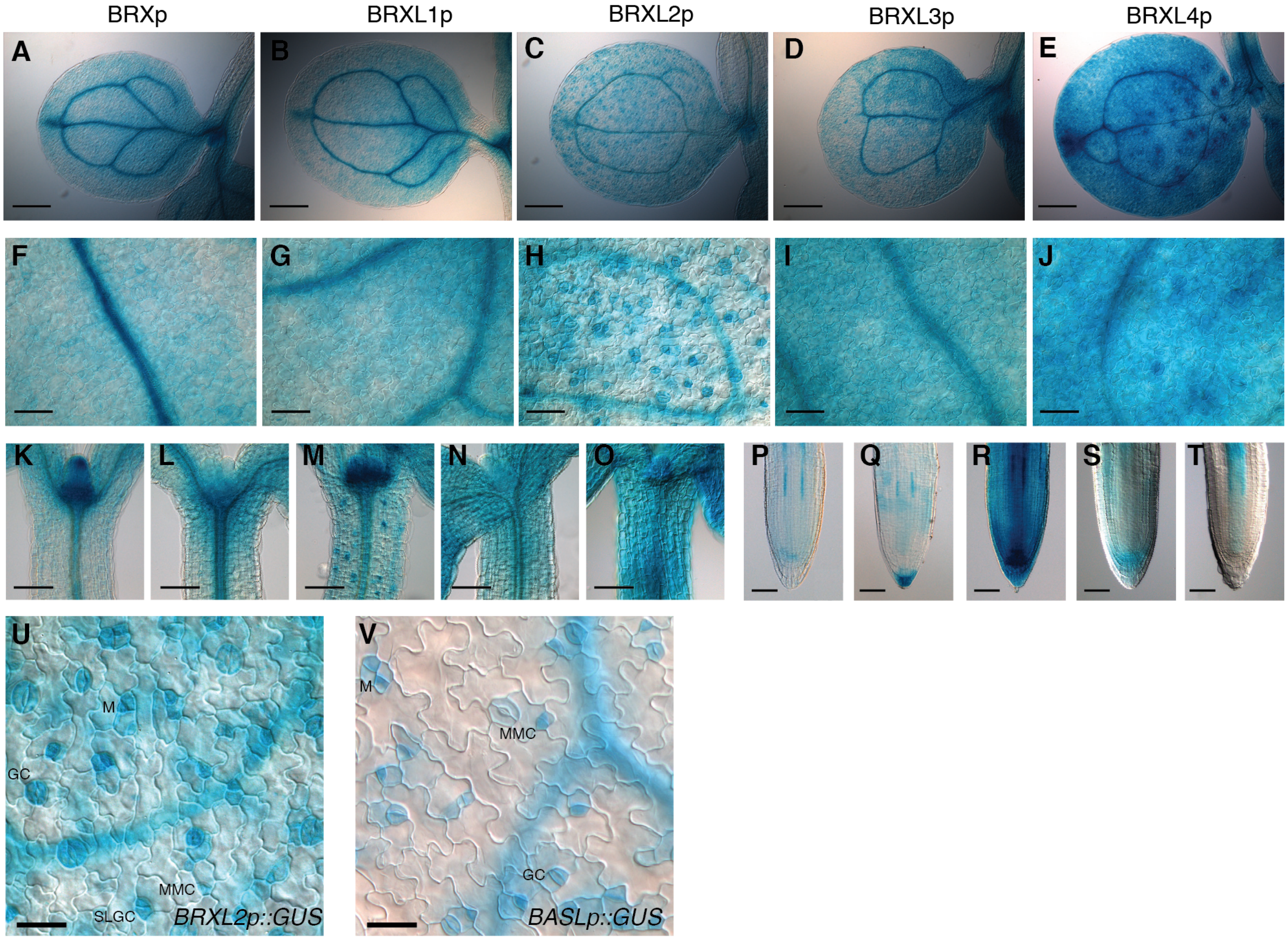


**Figure S2. The *BRX* gene family is expressed broadly in seedlings, with some members showing enrichment in the stomatal lineage**

**(A-U)** BRX family GUS transcriptional reporters in *Arabidopsis* seedlings at 3dpg. **(A-E)** Expression in abaxial cotyledons. The five reporters are listed in order at top of columns. **(F-J)** Close-ups of reporter expression in abaxial cotyledons. **(K-O)** Reporter expression in the hypocotyl and leaf primordia, order same as for A-E. **(P-T)** Reporter expression in primary roots, order same as for A-E. **(U)** Close-up of *BRXL2p:GUS* expression in cotyledon. Reporter expression in MMC, meristemoid, SLGC, and guard cells, respectively. **(V)** *BASLp:GUS* in cotyledon at 3dpg. Reporter expression in MMC, meristemoid, and guard cells, respectively. Scale bars: 200μm in (A-E), 50μm in (F-J, P-T), 100μm in (K-O), 30μm in (U-V).

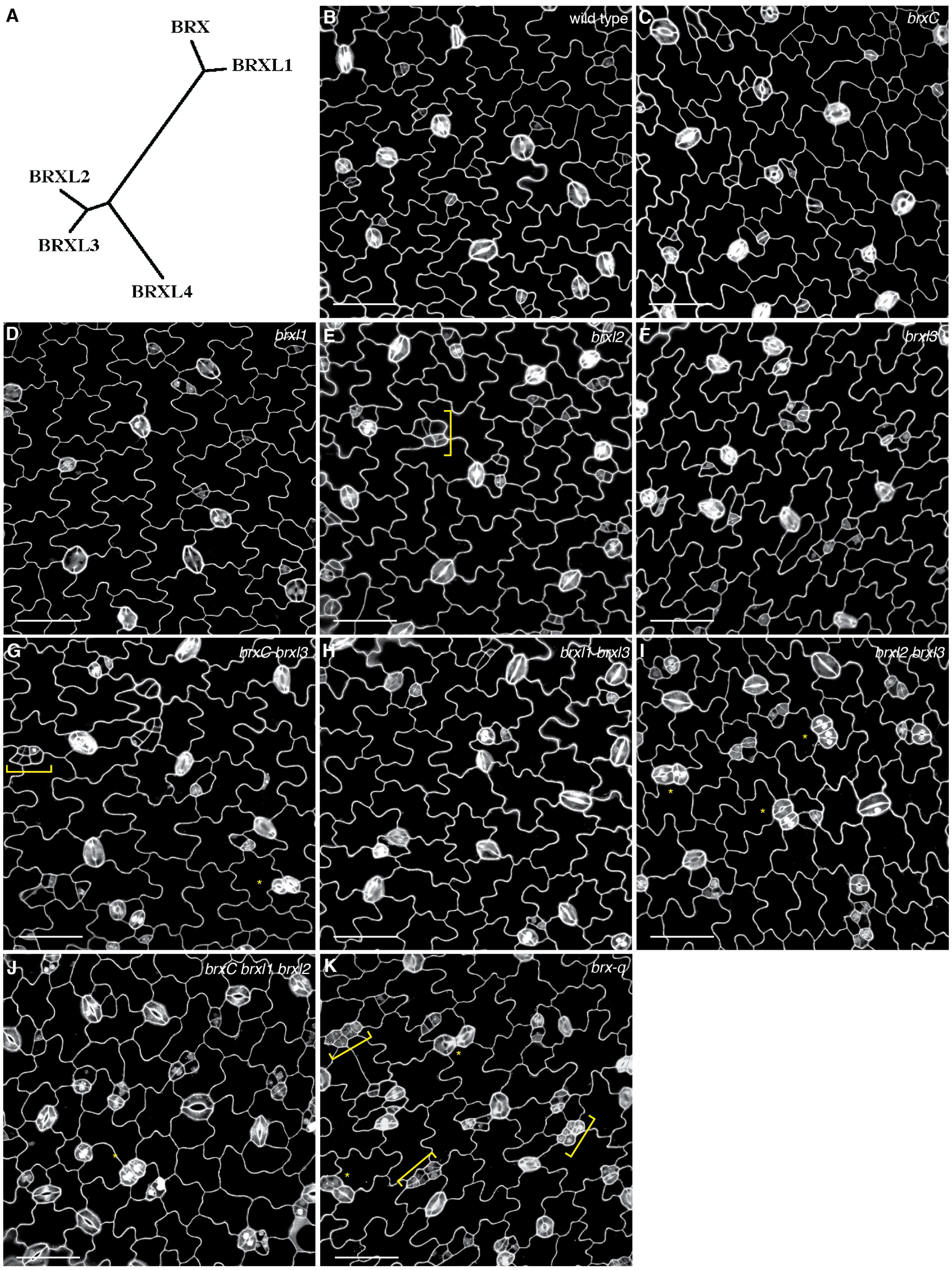


**Figure S3. BRX genes function redundantly in stomatal development**

**(A)** Schematic of relationship among Arabidopsis *BRX* gene family members (generated by maximum likelihood method based on amino acid sequences), line length indicates sequence distance. **(B-K)** Confocal images of 3dpg adaxial cotyledons, of single, double and triple *BRXf* mutant combinations. Genotypes indicated in panels. Cell outlines marked with propidium iodide. Yellow braces and asterisks indicate clustered small cells and stomata, respectively. Scale bars: 50μm.

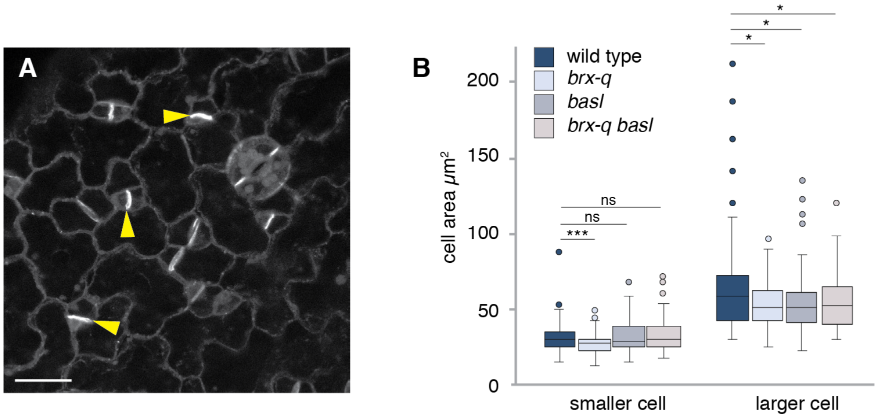


**Figure S4. Asymmetry defects are due to diminished size of larger daughter cells**
**(A)** Confocal image of 3dpg abaxial cotyledons stained with aniline blue. Arrowheads indicate callose-positive cell plates. Scale bar: 20μm. **(B)** Quantitation of cell areas for aniline-positive sister-cell pairs of 3dpg abaxial cotyledons. For each class, n = 100. * and *** show significant difference, P < .05, and P < .001, respectively by two-tailed Wilcoxon rank-sum test for the bracketed comparisons. Center lines in box plots show the medians; box limits indicate the 25th and 75th percentiles; whiskers extend 1.5 times the interquartile range from the 25th and 75th percentiles.

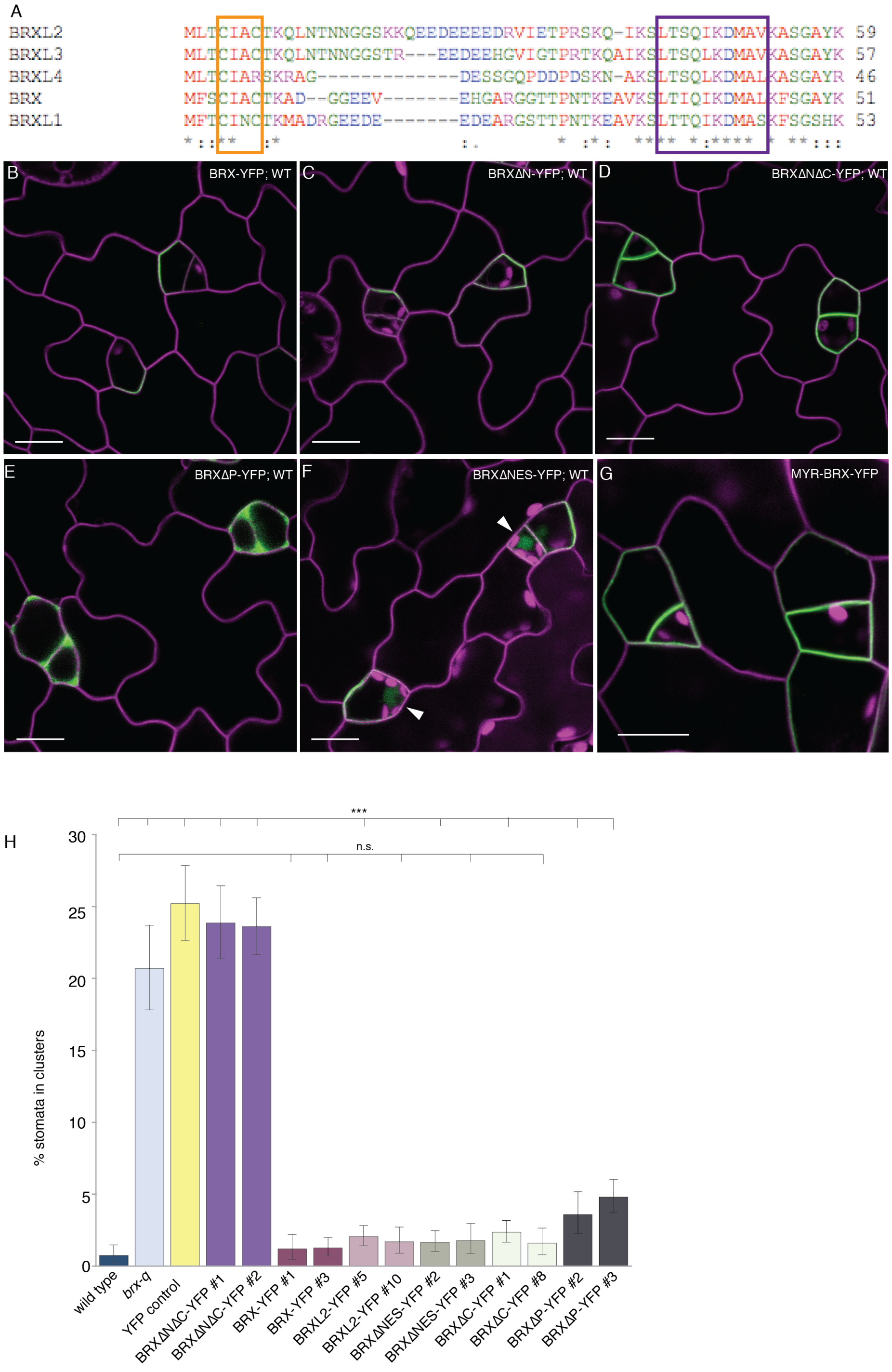


**Figure S5. Modifications to BRX protein sequence affect its localization**
**(A)** Alignment of N-terminal sequences of BRX family genes. Putative palmitoylation and nuclear export signals are boxed in orange and purple, respectively. **(B-G)** Confocal images of 3dpg abaxial cotyledons expressing YFP-tagged BRX proteins (green) and a PIP2A-RFP membrane marker (magenta). Genotypes as indicated on panels. Scale bars: 10μm. **(H)** Quantitation of rescue by various BRX constructs using stomata in contact in 7dpg adaxial cotyledons as an output. For each class, n = 20. Error bars represent SEM. Some lines in (H) are replicated here from Fig. 5H to enable a more convenient comparison among different lines.

**Figure S6. Supplemental tables quantifying stomatal clustering**
Detailed quantifications of stomatal cluster composition reported in Figures 2F, 5H and 5I.

**Supplemental Table for Fig 2F**

| | stomata | | | | | | |
|---|---|---|---|---|---|---|---|
| | single | in pairs | in triples | in quadruples | in quintuples | in sextuples | N |
| wild type | 836 | 0 | 0 | 0 | 0 | 0 | 836 |
| *brxl2 brxl2* | 729 | 184 | 18 | 8 | 0 | 0 | 939 |
| *brx-q* | 751 | 314 | 57 | 20 | 5 | 0 | 1147 |
| *basl* | 592 | 338 | 111 | 56 | 0 | 0 | 1097 |
| *brx-q basl* | 563 | 424 | 138 | 56 | 5 | 0 | 1186 |

**Supplemental Table for Fig 5H**

| | stomata | | | | | | |
|---|---|---|---|---|---|---|---|
| | single | in pairs | in triples | in quadruples | in quintuples | in sextuples | N |
| wild type | 805 | 10 | 0 | 0 | 0 | 0 | 815 |
| *brx-q* | 578 | 300 | 72 | 28 | 5 | 0 | 983 |
| *YFP control* | 832 | 400 | 72 | 8 | 0 | 0 | 1312 |
| *BRXΔNΔC-YFP* | 1349 | 630 | 225 | 84 | 15 | 12 | 2315 |
| *BRX-YFP* | 1238 | 30 | 0 | 0 | 0 | 0 | 1268 |
| *BRXL2-YFP* | 1566 | 60 | 6 | 0 | 0 | 0 | 1632 |
| *BRXΔNES-YFP* | 1238 | 40 | 0 | 0 | 0 | 0 | 1278 |
| *BRXΔC-YFP* | 1569 | 74 | 0 | 0 | 0 | 0 | 1643 |
| *BRXΔP-YFP* | 1265 | 108 | 27 | 0 | 0 | 0 | 1400 |

**Supplemental Table for Fig 5I**

| | stomata | | | | | | |
|---|---|---|---|---|---|---|---|
| | single | in pairs | in triples | in quadruples | in quintuples | in sextuples | N |
| wild type | 1108 | 16 | 0 | 0 | 0 | 0 | 1124 |
| *brx-q* | 1001 | 510 | 87 | 24 | 5 | 0 | 1627 |
| *YFP control* | 1127 | 562 | 96 | 28 | 0 | 0 | 1813 |
| *myrBRX-YFP L7* | 1045 | 110 | 12 | 0 | 0 | 0 | 1167 |
| *myrBRX-YFP L9* | 1020 | 66 | 0 | 0 | 0 | 0 | 1086 |

**Figure S7. Primer names and sequences used for cloning and genotyping**

| Purpose | Name | Sequence |
|---|---|---|
| BRX cDNA (Y2H vectors) | Brx cDNA Fw | CACCGAATTCATGTTTTCTTGCATAGCTTGTACC |
| (pGBK) | New Brx cDNA Rv | CGCGCTGCAGGAGGTACTGTGTTTGTATTCTCTCTC |
| (pGADT7) | Brx pGAD Rv III | CGCGGAGCTCGAGGTACTGTGTTTGTATTCTCTCTC |
| BRX-N (Y2H vectors) | BrxdomA Fw | CGCGGAATTCAAAGAGTGGATGGCTCAAGTAG |
| (pGBK) | BrxdomA Rv PstI | CGCGCTGCAGATTGTAAAGCTCAACGATCTTGTC |
| (pGADT7) | BrxdomA Rv SacI | CGCGGAGCTCATTGTAAAGCTCAACGATCTTGTC |
| BRX-C (Y2H vectors) | BrxdomB Fw | CGCGGAATTCGCTGAGTGGATTGAAGAGGA |
| | BrxdomB Rv | CGCGGGATCCGAGGTACTGTGTTTGTATTCTCTCTC |
| BRX promoter (pENTR) | At1g31880 5 Fw | CACCGTTTCAGCTGTCTCAAACCAATC |
| | At1g31880 5 Rv | GTCTCTTTTTTGAGTTGTTCTCTGCC |
| BRXL1 promoter (pENTR) | At2g35600 5 Fw | CACCCCTAGATCAAACTCACAGACTTTG |
| | At2g35600 5 Rv | GCACTCTTTTTTATGTTCTCTGCC |
| BRXL2 promoter (pENTR) | At3g14000 5 Fw | CACCCGAAACAGATCGTTGTGTAGAGTAC |
| | At3g14000 5 Rv | CTCACTAAAGAGTTTCAATTTCGACC |
| BRXL3 promoter (pENTR) | At1g54180 5 Fw | CACCGAGGGAATTTAAGGACAGATCG |
| | At1g54180 5 Rv | CTCTCACCTTCTCTTTTGACAC |
| BRXL4 promoter (pENTR) | At5g20540 Fw | CACCGGTTCTGCTAAAGGAGCAGATTC |
| | At5g20540 Rv | GACTGGCTTGGTCTTTGATTTGAG |
| *brx*$^{C}$ dCAPS genotyping | brx Fw (Uk-1) | CCATACCCTTTCATGGGTGGAAGT |
| | brx Rv (Uk-1) Hinf1 | GATATGAACACCAGGTTCTACTTGAGCGAT |
| *brxl1* (SALK_038885) genotyping | SALK_038885 LP | CGACTGAGCAGAGATGGATTC |
| | SALK_038885 RP | CAGACAGAGGTGAGGAGGATG |
| *brxl2* (SALK_032250) genotyping | SALK_032250 LP | TCAAAAGTTGACAAAATGCGG |
| | SALK_032250 RP | GCTTTTGTAAGCACCTGATGC |
| *brxl3* (SALK_017909) genotyping | SALK_017909 LP | TTCTGGGTTTTGCTTGAAATG |
| | SALK_017909 RP | GCCAAAATACCCATCCTTGAC |
| *basl-2* (WiscDsLox_264_F02) genotyping | basl-2 LP | ACAACTATCGGATCGTTGACG |
| | basl-2 RP | GAGGACGATTCCGTCTCTTTC |
| YFP cDNA (-stop) (pENTR NotI sites for N-terminal fusion) | YFP-NotI Fw (short) | CCGCGGCCGCATGGTGAGCAAGGGCGAGGAG |
| | YFP-NotI Rv | CCGCGGCCGCCTTGTACAGCTCGTCCATGCCGAG |
| BRXL2 promoter (pENTR NotI sites) | pBRXL2 NotI Fw | GCGCGGCCGCCGAAACAGATCGTTGTGTAGAGTAC |
| | pBRXL2 NotI Rv | GCGCGGCCGCCTCACTAAAGAGTTTCAATTTCGACC |
| BRXL2 cDNA (-stop) (pENTR) | BRXL2 Fw | CACCATGCTGACATGCATAGCTTGTACG |
| | BRXL2 Rv | CAAGTATTGTTGTTGTATCCGAGC |
| BASL promoter (pENTR NotI sites) | NewBASLpromoterNotI Fw | CCGCGGCCGCGGATATTCATCTTTGCTACACAAGAGTC |
| | NewBASLpromoterNotI Rv | CCGCGGCCGCGGCTGTTATGTTTGTGTTTCTTTGTCAC |
| BRX cDNA (-stop) (pENTR) | BRXcDNA Fw | CACCATGTTTTCTTGCATAGCTTGTACC |
| | BRXcDNA Rv | GAGGTACTGTGTTTGTATTCTCTCTC |
| BRX cDNA (-stop) (pENTR NotI/AscI sites) | OldSchoolBRXintopENTR Fw | CCGCGGCCGCCATGTTTTCTTGCATAGCTTGTACC |
| | OldSchoolBRXintopENTR Rv | CCGGCGCGCCCGAGGTACTGTGTTTGTATTCTCTCTC |

| | | |
|---|---|---|
| BRX promoter (pENTR NotI sites) | pBRX-NotI Fw | CCGCGGCCGCGTTTCAGCTGTCTCAAACCAATC |
| | pBRX-NotI Rv | CCGCGGCCGCGTCTCTTTTTTGAGTTGTTCTCTGCC |
| BRX∆C cDNA (PCR mutagenesis of BRX) | BRX (minus C-terminal BRX domain) Rv | TTGCATTTCACTAGCATTGCTCATG |
| BRXΔN cDNA (PCR mutagenesis of BRX) | Fw (5 of N-terminal BRX) | GTACTGGATGATGATGGACCAGTACAGAGATTTAACCGCCAAG |
| | Rv (3 of N-terminal BRX) | CTTGGCGGTTAAATCTCTGTACTGGTCCATCATCATCCAGTAC |
| BRXC$^{4}$C$'$→A$^{4}$A$'$ (BRXΔP) (PCR mutagenesis of BRX) | BRXminusP Fw | CACCATGTTTTCTGCCATAGCTGCTACCAAAGCAG |
| BRXΔNES cDNA (PCR mutagenesis of BRX, substitution of NES residues) | BRXNEStoA Fw | AAAAGCGCAACCATTCAGGCTAAAGATGCGGCTGCGAAATTC |
| | BRXNEStoA Rv | GAATTTCGCAGCCGCATCTTTAGCCTGAATGGTTGCGCTTTT |
| BRXΔNES cDNA (PCR mutagenesis of BRX, deletion of NES) | BRXdNES Fw | ATAAGCACCAGAGAATTTGCTTTTGACGGCTT CTTTAGTATTGGGAGT |
| | BRXdNES Rv | AAGAAGCCGTCAAAAGCAAATTCTCTGGTGCTTATAAACAATGCAAGCA |
| MYR-BRX cDNA (addition of N-terminal myristoylation signal by PCR) | Myr-BRX Fw | CACCATGGGCAACAAATGTTGCAGCAAGCGACAGGATACCATGGCCAT GTTTTCTTGCATAGC TTGTACCAAAGC |
